# A heuristic underlies the search for relief in fruit flies

**DOI:** 10.1101/2020.06.01.127753

**Authors:** Nicola Meda, Giulio Maria Menti, Aram Megighian, Mauro Agostino Zordan

## Abstract

Humans rely on multiple systems of sensory information to make decisions. However, strategies that shorten decision-making time by taking into account fewer but more essential elements of information are preferred to strategies involving complex analyses. These “shortcuts to decision” are also termed “heuristics”. The identification of heuristic principles in species phylogenetically distant to humans would shed light on the evolutionary origin of speed-accuracy trade-offs and offer the possibility to investigate the brain representations of such trade-offs, urgency, and uncertainty. During experiments on spatial learning, we acknowledged that the search strategies of the invertebrate *Drosophila melanogaster*, the common fruit fly, resembled a spatial heuristic. Here we show that the fruit fly applies a heuristic termed the “Nearest Neighbour Rule” to avoid bitter taste (a negative stimulation). That is, the fly visits the salient location closest to its current position to hopefully stop the negative stimulation. Only if this strategy proves unsuccessful, the animal uses other learned associations to avoid bitter taste. The acknowledgement of a heuristic in *D. melanogaster* supports the view that invertebrates can leverage on ‘economic’ principles when making choices and that the existence of heuristics in evolution dates to at least 600 million years ago.

## Introduction

Navigating an environment is a complex process, and several organisms across different *taxa* have been studied to address how animals apply diverse strategies to move in the surrounding environment ^1,2^. The picture that emerges from these studies is that different animals can apply analogous behaviours during similar navigation demands to reach their targets ^3–5^, such as food, shelter, or peers. To achieve their goals, animals frequently have to make adequate decisions by leveraging on different sources of information ^6,7^. However, considering multiple cues before making a choice is an effortful strategy that may hinder the chances of survival ^8^. Instead of a piecemeal, in-depth analysis of sensory information, humans appear to privilege strategies that minimise cognitive load (i.e., use the least amount of information) and shorten decision-making time, but still lead to appropriate and adaptive behaviour ^9,10^. These strategies are also named “heuristics” ^11^. For example, option discrimination can rely on a heuristic termed “Take The Best”. According to this heuristic, only the most reliable cue in terms of discriminatory capacity between different outcomes ^12^ (e.g., punishment or reward) is used for inference, while cues with lower predictive value are not taken into consideration ^13,14^. Several sub-optimal choices made by nonhuman animals could be deemed to be based on heuristic principles, yet such behaviours are rarely adequately acknowledged ^15^. Given that heuristics are “shortcuts” to decisions, animals are expected to implement these strategies under uncertainty ^16^, during highly complex tasks ^17^, or in urgency. Only later, environmental conditions allowing, or if the goal has not been achieved, animals may adjust their decisions accordingly ^18^. Moreover, diverse spatial heuristics (strategies applied specifically during navigation) have been described in humans, thus a comparative search for the same strategies in other animals could be relatively straightforward ^17,19^. One of these heuristics is the “Nearest Neighbour Rule” (NNR), or proximity rule. According to NNR, a moving animal searching for food, or other relevant stimuli, visits the closest location to its current position first, then travels to the next closest position, and so on ^20^. Most of the research on the subject has been conducted in the field of behavioural economics, on human participants ^21^. Thus, there is a knowledge gap in the origin of heuristic principles and their conservation through evolutionary time, whereas this information could represent a fruitful entry-point to investigate uncertainty and urgency in nonhuman animals. This lag is understandable considering the complexity of adapting tasks specifically designed for humans to other animals. Nonetheless, we argue that short-range navigation tasks (i.e., conducted in a laboratory) satisfy the criteria for investigating heuristic principles from a comparative standpoint: animals can be forced to choose under uncertainty – by controlling the explorable environment ^22^, in urgency – by stimulating them negatively, or by designing more complex tasks ^23^.

Identifying heuristic principles in phylogenetically distant species would shed light on the origin of speed-accuracy trade-offs ^24,25^. The recognition of the same principles in model organisms would represent the entry-point to investigate the brain representations of such trade-offs, of “cognitive load”, of urgency and uncertainty.

In this study on spatial learning, we show that fruit flies are able to associate relief from bitter taste (a negative stimulation that is innately associated with a threat of intoxication) to a specific visual landmark and search for the expected outcome (i.e., relief) in the proximity of the landmark. However, we acknowledged that the initial search strategies of the invertebrate *Drosophila melanogaster* resemble a spatial heuristic. Specifically, we observed that in order to relieve an unpleasant stimulation, fruit flies first performed a search near their current position and shortly after approached the appropriate landmark. We thus conducted a set of experiments aimed at investigating whether this behaviour was due to insufficient learning or whether it was the result of a trial-and-error strategy. Our data suggest instead that the strategy of fruit flies is in line with that of pigeons ^26^, primates ^24^, rats, and humans ^27^, and that the initial search behaviour of the flies can be explained by the application of the Nearest Neighbour Rule, whereas learned visual information is used only if the former strategy is unsuccessful.

## Results

*Drosophila melanogaster* can be trained to distinguish identical objects with different orientations ^28–30^. A blinded experimenter used a circular arena to train single flies (n = 40) to differentiate between a black vertical stripe, which marked the presence of a “safe zone” (6 cm^2^ surface) linked to relief from optogenetically-induced bitter taste, from a diametrically-opposed horizontal stripe which was not associated to relief. The training session was composed of 16 trials (i.e., repetitions), each three minutes long. During the first 30 seconds of each trial, the fly was free to explore the arena in complete darkness; for the next 30 seconds, the fly could explore the arena in the presence of the diametrically opposed visual patterns (black horizontal/vertical bar on a homogenously-lit background). In the last 2 minutes, the fly could experience bitter taste according to its position in the arena, while still in the presence of the visual patterns. At the beginning of each new trial, the positions of the matched vertical stripe – safe zone and of the horizontal stripe were switched (Figure 1A. Further details regarding materials and methods are available as supplementary materials). We considered the time spent in the safe zone throughout the training session (Figure 1B, Table S3), and the preference for the vertical bar (measured in terms of time spent in its proximity) during the probe session (Figure 1E) as a proxy for learning (Figure S1-S2, Table S1-S2 for control experiments). At the start of each new trial, given the switch of the positions between the horizontal bar and vertical bar-safe zone, fruit flies could experience two potentially conflicting sources of information as to the current spatial location of the safe zone (CSZ). The CSZ could be located either in the spatial location where relief was last experienced during the preceding trial (where the horizontal stripe is currently displayed) or in the proximity of the vertical bar. In this cue-conflict paradigm, the animal could employ a local search strategy centred on the spatial location most recently associated with relief or use the learned visual association between the vertical stripe and relief to approach the safe zone. The analysis of data was conducted by considering only the trials during which each fly was not already inside one of the two areas of interest, i.e., the current safe zone (CSZ) or the previous safe zone (PSZ), at the onset of stimulation.

**Fig. 1.**
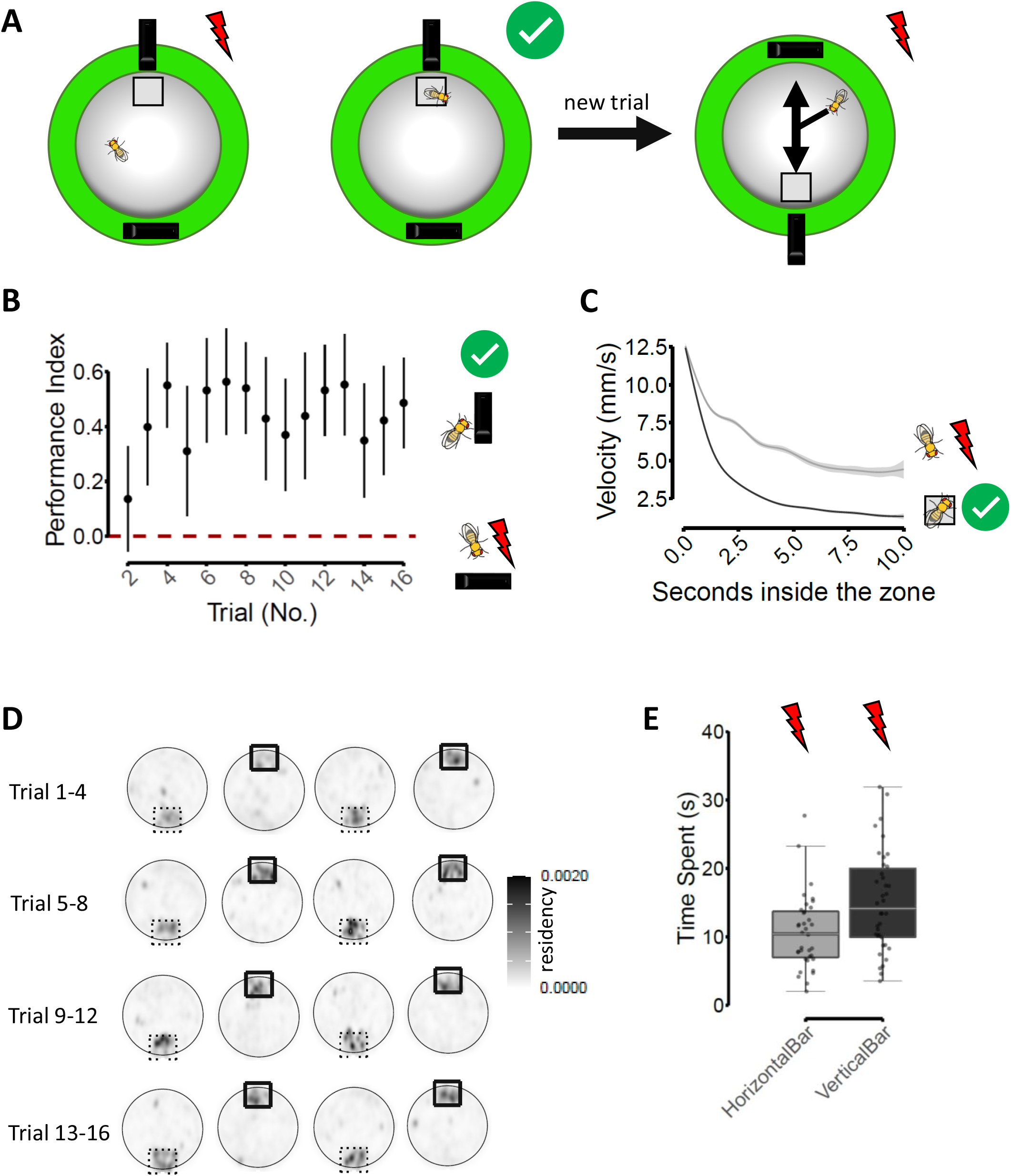
Behavioural paradigm and fruit flies learning. **A)** summary of the behavioural paradigm and experimental question. Bitter stimulation depends on the fly’s position in the arena. At the beginning of a new trial, does the fly resort to the Nearest Neighbour Rule to stop the bitter stimulation, or will it approach the visual marker? **B)** Performance Index (PI) as a function of training trials. PI = the difference between the time spent in the safe zone and the time spent in the previous (location of the) safe zone, divided by the total time spent in the two zones. The performance index can range from 1 to -1. A value of zero indicates no zone preference. PI was not computed for trial 1 given the absence of a Previous Safe Zone during that trial. Pointrange = mean ± confidence interval around the mean. Flies spend more time in the safe zone with respect to the previous safe zone, marked by the vertical stripe, throughout training (see also Table S3); **C)** Velocity profile of fruit flies after entering a zone where relief is provided (black line) compared to the profile (grey line) after entering the non-safe zone (Table S4); **D)** density plots describing the residency of flies during the training session, when bitter-taste stimulation is triggered if flies leave the safe zone (squared); **E)** During the probe session, the vertical bar was located either at the northern, southern, eastern or western end of the arena (n = 10 flies were tested for each location). Fruit flies spend more time near the vertical stripe than near the horizontal one even though relief from bitter-taste cannot be achieved in any zone (no. observations = 75, VerticalBar - HorizontalBar mean difference in time spent (s) 5.03, std error = 1.47, z.ratio = 3.14, p = 0.0017).

We found that flies significantly increased the number of visits to both zones (PSZ and CSZ) during the first 10 seconds of optogenetic stimulation (Figure 2A, Table S4, no. observed visits = 151, model estimate mean (log-scale) = 0.883, std error = 0.15, z.ratio = 5.61, p < 0.0001) with no preference for either of the two zones. This increase is also reflected by the higher mean number of flies that can be found in both zones during this period of time, with respect to what was observed 10 seconds before the onset of stimulation (Figure 2B, Table S4, no. observed flies = 89, model estimate mean (log-scale) difference = 0.80, std error = 0.12, z.ratio = 6.619, p < 0.0001). The fact that the PSZ and the CSZ were visited in the same measure by the same number of flies suggests that flies did not take into consideration which visual marker they were approaching (although they are able to learn to differentiate between two identical shapes with different orientations, Figure 1 and ref. ^28–30^). Thus, we hypothesised that the animals first tend to direct themselves to the zone closest to their current position (a rapid behaviour), and only later do they appear to take into consideration the orientation of the visual marker (a slower but more accurate response) ^14,31^. In order to evaluate whether this was the case, we proceeded to identify the positions of flies at the onset of optogenetic stimulation and separated the animals into two groups according to which was the first zone entered. Figures 2C and 2F show the positions of the flies which entered the CSZ or the PSZ (marked by the horizontal bar) respectively, during the trials in which the CSZ was located at the “northern” end of the arena (data on trials in which the CSZ was located at the “southern” end are presented, for ease of readability, in Figure S3). These density plots suggest that flies tended to enter the zone closest to them at the onset of the optogenetic stimulation, a behaviour consistent with the use of the Nearest Neighbour Rule. Next, we tested whether the positions of each group of flies for each set of trials (i.e., subdivided according to whether the CSZ was at the northern or southern end of the arena) were more aggregated than expected under the null hypothesis of a random distribution. To do this, we applied Marcon and Puech’s M function ^32–34^ and tested if the observed positions of the flies were more aggregated than expected based on the generation of 10.000 simulated random positions. For both sets of trials, and for both groups of flies (i.e., the flies grouped according to which zone was entered first), animals showed evidence for significantly greater aggregation than expected (Figure 2D goodness-of-fit test testing for aggregation p = 0.012 (meaning M(r) > 1), for n = 63 positions tested, Figure 2G goodness-of-fit test p = 0.030, for n = 62 positions, and Figure S3). Moreover, if the position of a fly in the arena is a reasonably good predictor of which will be the first zone visited, the positions of the flies which are predicted to enter the PSZ should be spatially segregated from the positions of the flies that are predicted to enter the CSZ. In fact, we found that the positions of the two groups of flies show significant segregation in both sets of trials (Figure 2J goodness-of-fit test testing for spatial repulsion p = 0.031 (meaning M(r) < 1) for n = 125 positions, and Figure S3).

**Fig. 2.**
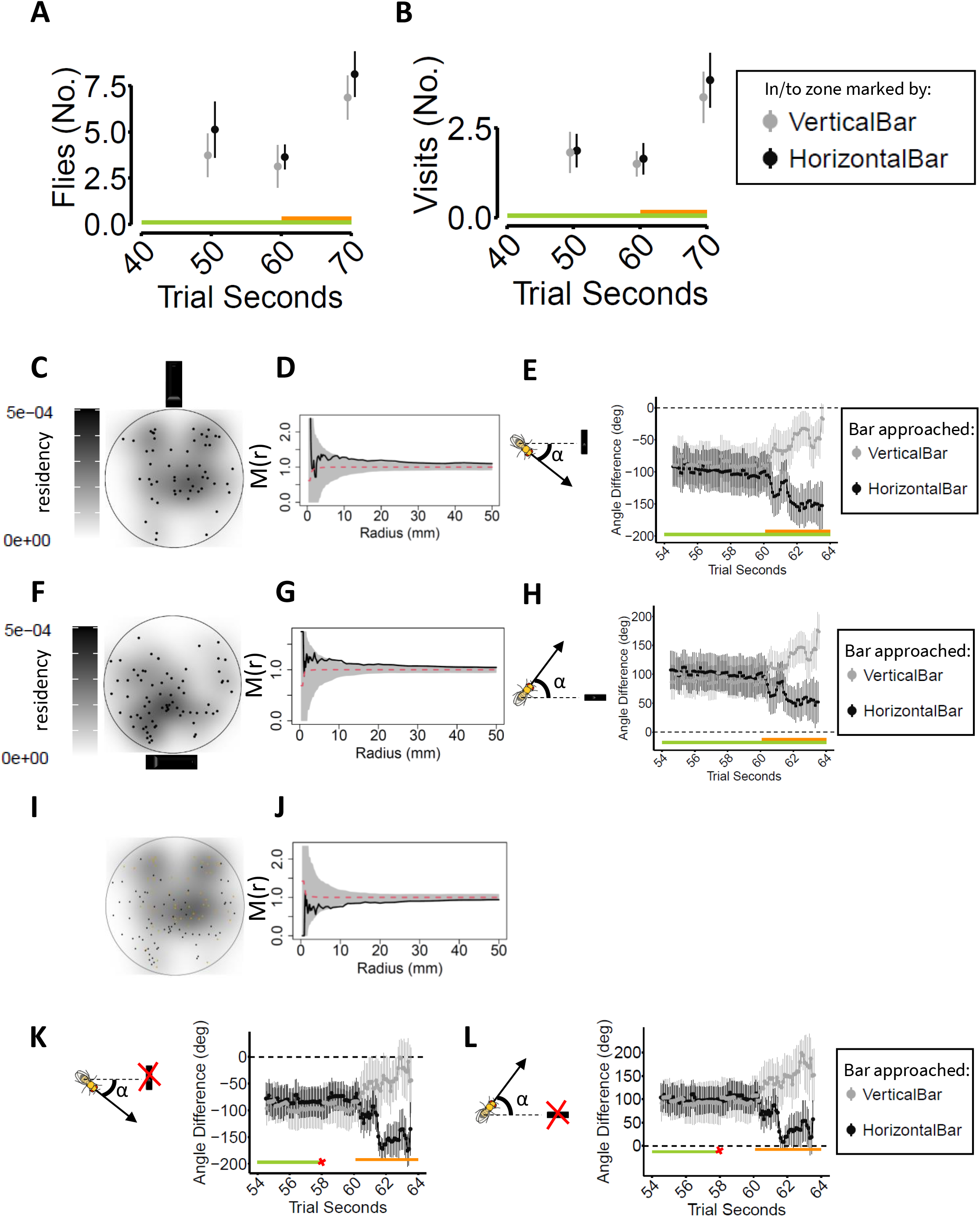
Fruit flies resort to the Nearest Neighbour Rule soon after bitter stimulation onset. Even-numbered trials. Green bar = visual patterns displayed; orange bar = stimulation triggered according to fly position. Pointrange = mean ± confidence interval around the mean. **A)** In the 10 seconds after the onset of optogenetic stimulation, the difference between the number of flies that approached either one of the two zones is not significant (Table S4); **B)** There is no difference between the mean number of visits to the two zones in the periods considered, further implying that flies searched both zones in the same measure (Table S4); **C)** Spatial position of fruit flies, at second 60.1, (when the first pulse of optogenetic stimulation is delivered) that first entered the safe zone, marked by the vertical stripe **D)** Marcon and Puech’s M function value (black line) represents the distance between the observed positions of flies compared to 10.000 random distribution simulations (red dashed line and grey shading). A value greater than 1 suggests aggregation. A goodness-of-fit test reveals that flies are significantly more aggregated than expected; **E)** fruit flies that entered the safe zone oriented themselves towards that zone after the onset of the bitter stimulation; **F)** Spatial distribution at second 60.1 of fruit flies that will approach the previous safe zone; **G)** M function value is significantly greater than expected under the null hypothesis, suggesting aggregation of flies; **H)** the fruit flies that entered the previous safe zone oriented themselves towards it (see also Table S4); **I)** Spatial position of both groups of flies at second 60.1; **J)** M function value assessing whether the two distributions of flies’ positions reported in C), F) or I) consist of two distinct aggregates. The M function is < 1, thus suggesting spatial repulsion between the two groups of flies, indicating that flies entering the safe zone are spatially segregated from flies that enter the previous safe zone. **K)** the fruit flies re-orient themselves towards the expected location closest to the landmark (in this case the vertical stripe, even though the landmark is occluded from vision), suggesting that flies retrieve the position of the visual target from working memory. **L)** as for K), in this case the horizontal stripe would be the closest landmark that is occluded from vision.

Furthermore, if the observed behaviour of the flies is truly the result of the application of a Nearest Neighbour (NN) model, the flies should not enter a given zone by chance, because in this case, the behaviour would be better explained by random navigation ^19^. In other words, in the NN model, the animals should first orient themselves towards the closest zone, irrespective of the visual marker associated with it, and then travel towards it. Our data show that, shortly after the beginning of the optogenetic stimulation, fruit flies re-orient themselves towards the closest zone before approaching it (for flies approaching the vertical bar, Figure 2E, no. orientations tested = 8937, mean orientation difference after-before the onset of stimulation, in degrees = 25.5, std error = 2.35, z.ratio = 10.84, p < 0.0001; for flies approaching the horizontal bar, Figure 2H mean orientation difference = 36.2, std error = 2.14, z.ratio = 16.86, p < 0.0001, and Figure S3). Moreover, the target towards which the fly orients itself should depend on the position of the fly in the arena. If this is indeed the case, the orientation of the animal at the onset of stimulation (second 60.1), after controlling for the position, should not be a good predictor of the zone which will then be approached. We thus modelled the zone that each animal would eventually enter based on the animal’s position alone or based on both position and orientation (Table S4). We found that the fly position in the arena is necessary and sufficient to predict which zone the fly will enter first, and the addition of orientation to the model does not provide a better explanation of the observed data. Lastly, we tested other flies (n = 30) under a different paradigm: during each trial, 2 seconds before the onset of optogenetic stimulation, and 4 after the onset, we presented the flies solely the homogeneously lit background, without landmarks (thus the stripes were present until second 58 of the trial and reappeared after second 64). We ascertained that flies re-oriented themselves towards the supposed location of the landmark and travelled towards the expected position, although no stripe was present (Figure 2K, no. orientations tested = 6917, mean orientation difference after-before the onset of stimulation, in degrees = 44.3, std error = 3.09, z.ratio = 14.36, p < 0.0001; Figure 2L, mean orientation difference = 37.2, std error = 2.4, z.ratio = 15.45, p < 0.0001. see Figure S3 for odd-numbered trials). As previously reported, this behaviour can be explained by considering that the animal is using the memory of the landmark position in the environment to guide its goal-directed navigation ^35^. We conducted a replicate set of the experiments with a new group of flies (n = 41), further supporting the results presented here (Figure S4, Table S5-S6). We also conducted an additional set of experiments with 40 more flies, in which the animals were tested in a visual environment with two identical and diametrically opposed vertical bars. We alternated the position of the safe zone between the two bars on different trials. With this set of experiments, we extend the results suggesting that the initial search strategy (resorting to the Nearest Neighbour Rule) employed by the flies appears to be independent of the width or orientation of the visual cues (Figure S5, Table S7-S8). These experiments further confirmed that the zone first entered by the fly is predicted by the spatial location of the animal at the onset of the negative stimulation, thus supporting that this behaviour resembles the spatial heuristic known as the Nearest Neighbour Rule ^19,23,36,37^.

## Discussion

In order to survive, animals have to evaluate environmental and internal information and make decisions rapidly. However, rapidity comes with imprecision, and the trade-off between speed and accuracy differentiates between a perfect but inapplicable strategy, a fast and frugal strategy, and an immediate but unsuccessful approach ^8,14,31^. Heuristics are shortcuts extensively used by humans, characterised by an optimal balance between speed and accuracy ^37–39^. The use of these shortcuts can be elicited by pressing the animal with urgency, uncertainty, or cognitive load (e.g., by forcing it to accomplish two tasks simultaneously) ^14,17^. In contrast to the analysis of heuristics in human cognitive psychology and behavioural economics, the investigation of these strategies in behavioural neuroscience has lagged behind, possibly due to the inapplicability of the economic paradigms in animals other than humans, but probably also because heuristics, when unacknowledged, are regarded as behavioural “noise” or as being due to insufficient learning ^15^. During experiments on spatial learning, conceptually similar to the paradigms described in ^22,36^, we showed that fruit flies learned the spatial association between a vertical bar and relief from an unpleasant stimulation produced via optogenetic stimulation of bitter-sensing neurons. However, we also acknowledged that the search behaviour of *Drosophila melanogaster* at the onset of the stimulation resembled a spatial heuristic. We fortuitously elicited this spatial heuristic probably because we exposed the flies to bitter taste, which is a negative but ecologically relevant stimulation (thus providing urgency) that nonetheless can be leveraged in a learning paradigm ^40^. Moreover, under these conditions, flies faced conflicting cues regarding where the relief from the stimulation could take place (uncertainty). Urgency and uncertainty are two circumstances that trigger the emergence of heuristic behaviour ^9,38^. With this paradigm, we observed that fruit flies initially adopt the heuristic known as the “Nearest Neighbour Rule” (NNR) to escape punishment, and only if this proves unsuccessful, they resort to other strategies. According to the NNR, a moving animal seeking food or ecologically relevant information should visit the closest location to its current position first, and then travel to the next closest position, and so on ^20^. In our experiments, the fruit flies approached the closest visual marker to their current position, or the closest position where they expect the landmark to be (even in the absence of a landmark, Figure 2K and 2L ^35^). Only if this goal-oriented navigation proved unsuccessful, the animals resorted to the learned visual information in order to locate the safe zone. This heuristic was already described in mammals ^37^ as well as other vertebrates ^23^, but the evidence herein supports the idea ^25^ that also insects employ “shortcuts” to decision, and that from an evolutionary point of view the existence of heuristics, in particular of the NNR, might date to the last common ancestor of Arthropods and Vertebrates ^41^.

## Acknowledgements

The authors thank Dr. Paola Cisotto for technical support, Dr. Giovanni Frighetto for experimental set-up preparation, Valentina Tanara (blinded experimenter) for support with data acquisition, Prof. Marco Dal Maschio and Dr. Irene Slongo and Dr. Alberto Bettella for helpful suggestions, and Prof. Christian Wegener for providing BSC line 55135.

## Author contributions

Conceptualization: NM, GMM, AM, MAZ

Methodology: NM, GMM, AM, MAZ

Software: MAZ

Investigation: NM, GMM

Formal analysis: NM

Visualization: NM

Funding acquisition: AM, MAZ

Supervision: AM, MAZ

Writing – original draft: NM

Writing – review & editing: GMM, AM, MAZ

## Data and materials availability

Datasets are available at https://data.mendeley.com/datasets/9frwpy5vz9/draft?a=5e9ec4cc-7e52-44d7-af0d-38718078de0f (definitive DOI will be provided after review), customised MatLab script at ^42^.

